# Contrasting signatures of interspecific gene flow in the Atlantic Forest endemic butterfly *Heliconius besckei*

**DOI:** 10.64898/2026.10.05.756708

**Authors:** Mar Repullés, Thibaut Capblancq, Jan Kollross, Kay Lucek, Mathieu Joron, Niklas Wahlberg, Karina Lucas Silva-Brandão, André V.L. Freitas, Pável Matos-Maraví

**Affiliations:** Biology Centre CAS, Institute of Entomology, České Budějovice, Czech Republic; Faculty of Science, University of South Bohemia, České Budějovice, Czech Republic; LECA, Université Grenoble Alpes, Université Savoie Mont-Blanc, CNRS, Grenoble, France; University of Neuchâtel, Neuchâtel, Neuchâtel, Switzerland; CNRS, Centre d’Ecologie Fonctionnelle et Evolutive, Université de Montpellier, France; Department of Biology, Lund University, Lund, Sweden; Museum of Nature Hamburg Zoology, Leibniz Institute for the Analysis of Biodiversity Change, Hamburg, Germany; State University of Campinas UNICAMP, São Paulo, Brazil

**Keywords:** speciation with gene flow, *Heliconius ethilla narcaea*, *Heliconius melpomene nanna*, demographic models, species distribution model, whole-genome sequencing

## Abstract

Gene flow between diverging species can reshuffle genomic variation and greatly influence the speciation process, sometimes reinforcing, sometimes eroding species boundaries. However, the effect of ecological and historical processes on the patterns of introgression remains poorly understood. Here, we focus on the evolutionary history of *Heliconius besckei*, a geographically restricted butterfly endemic to the Brazilian Atlantic Forest, together with the mimetic *H. melpomene nanna*, the sympatric non-mimetic *H. ethilla narcaea*, and its closely related *H. numata robigus*. Using whole-genome sequencing, genome-wide tests of introgression, demographic modelling and palaeoclimatic species distribution models, we tested the extent and timing of interspecific gene flow in relation to mimicry, ecological context, and changes in geographic overlap. Despite strong genomic differentiation among the four focal *Heliconius* species, we detected widespread signatures of interspecific gene flow, greatly influenced by geographical overlap. Demographic models including post-divergence gene flow were better supported than strict-isolation models. In addition, genome-wide introgression analyses showed more numerous and larger candidate introgressed regions between *H. besckei* and *H. ethilla narcaea*, consistent with recurrent opportunities for exchange between these currently sympatric species. Species distribution models further recovered repeated expansions of suitable habitat for *H. besckei* during cold periods, providing a potential geographic context for secondary contact within the Atlantic Forest. Together, our results reveal recurrent gene flow occurring under different temporal, geographic, and potentially selective contexts, and show that the evolutionary history of a geographically restricted lineage can be shaped by repeated genetic exchange while substantial species differentiation is maintained.

## 1. Introduction

Gene flow between species is widespread across the Tree of Life, often contributing to the evolution of adaptive traits and shaping genome-wide variation across species (Mallet 2005; The Heliconius Genome Consortium 2012; Abbott et al. 2013). Increasing genomic evidence suggests that species diversification frequently occurs despite ongoing gene flow, resulting in complex evolutionary histories and heterogeneous patterns of introgression along the genome (Abbott et al. 2013; Moran et al. 2021). Rather than only eroding species boundaries, interspecific gene flow can facilitate adaptation through the transfer of beneficial alleles among diverging lineages (Moran et al. 2021; Leroy and Heuertz 2026) and promote species diversification through the creation of new genetic combinations (Seehausen 2004; Capblancq et al. 2015; Rosser et al. 2024). However, the factors determining where, when, and to what extent gene flow occurs between diverging species remain poorly understood.

Butterflies of the genus *Heliconius* (Lepidoptera: Nymphalidae) provide a great system for investigating the dynamics of interspecific gene flow, as extensive hybridisation can occur despite persistent species boundaries (Brower 2012; Kozak et al. 2021; Cicconardi et al. 2023; Thawornwattana et al. 2023). One of the best documented examples involves the evolution of their wing colour patterns. Multiple *Heliconius* species converge on shared warning colours through Müllerian mimicry, often facilitated by adaptive introgression of wing patterning alleles (Sheppard et al. 1985; Mallet and Gilbert 1995; Salazar et al. 2010; Hines et al. 2011; Reed et al. 2011; Pardo-Díaz et al. 2012; The Heliconius Genome Consortium 2012; Wallbank et al. 2016; Mérot et al. 2013). Much of this phenotypic variation is regulated by a small set of major-effect loci, including regulatory regions associated with the genes *optix*, *cortex* and *WntA* (Reed et al. 2011; Kronforst and Papa 2015; Nadeau et al. 2016), as well as the supergene *P* in *H. numata* (Joron et al. 2011). Moreover, genomic studies increasingly indicate that introgression in *Heliconius* is not restricted to mimicry loci and can involve substantial portions of the genome, with ecological and geographic context influencing patterns of gene exchange (Montejo-Kovacevich et al. 2022). Thus, *Heliconius* provides the opportunity to study how ecological interactions and historical changes in species distributions shape the direction and extent of interspecific gene flow.

Most genomic studies of *Heliconius* have focused on Amazonian and Andean lineages, where species diversity and mimicry are particularly high (Merrill et al. 2015; Jiggins 2017). In contrast, much less is known about the evolutionary dynamics of the Atlantic Forest lineages in eastern South America. The Atlantic Forest has a distinct biogeographic history shaped by past connections with the Andes and Amazon forest, complex topography and climatic fluctuations, resulting in a pronounced geographic structure of populations and species, as well as high endemism (Carnaval & Moritz 2008; Peres et al. 2020; Repullés et al. 2026). In particular, the southeastern mountain ranges have been proposed as relatively stable refugial regions that facilitated lineage persistence while climatic oscillations repeatedly altered the extent and connectivity of suitable habitats (Brown 1979; Carnaval et al. 2009; Pazt et al. 2018). Thus, the Atlantic Forest provides an ideal natural system for investigating whether pervasive gene flow documented throughout the *Heliconius* radiation has also shaped the diversification of geographically restricted and historically isolated lineages.

*Heliconius besckei* is mostly restricted to the mountains of the Atlantic Forest of southeastern Brazil (Brown & Mielke 1972). Phylogenomic analyses place *H. besckei* in an early-diverging lineage of the silvaniform assemblage, a *Heliconius* clade characterised by tiger-like wing patterns resembling Ithomiini butterflies, with the *H. numata*-*H. ismenius* lineage as its closest extant relatives (Zhang et al. 2016; Kozak et al. 2021). These three species diverged from the remaining silvaniform–melpomene clade approximately 3–4.8 Mya (Cicconardi et al. 2023; Thawornwattana et al. 2023). Unlike most members of the silvaniform assemblage, *H. besckei* exhibits a *postman* pattern consisting of a black forewing with a transverse red band and a yellow hindwing bar (Figure 1A). In the Atlantic Forest, this pattern is shared with *H. melpomene nanna* and *H. erato phyllis*, whereas *H. melpomene burchelli* occurs further west along the Cerrado biome region (Brown 1979, Rosser et al. 2012; Rosser & Mallet 2024). The red and yellow components of the *postman* phenotype are linked to the genes *optix* and *cortex*, whose expression is controlled by the B/D and the Yb regulatory regions, respectively (Reed et al. 2011; Nadeau et al. 2016; Wallbank et al. 2016).

**Figure 1.**
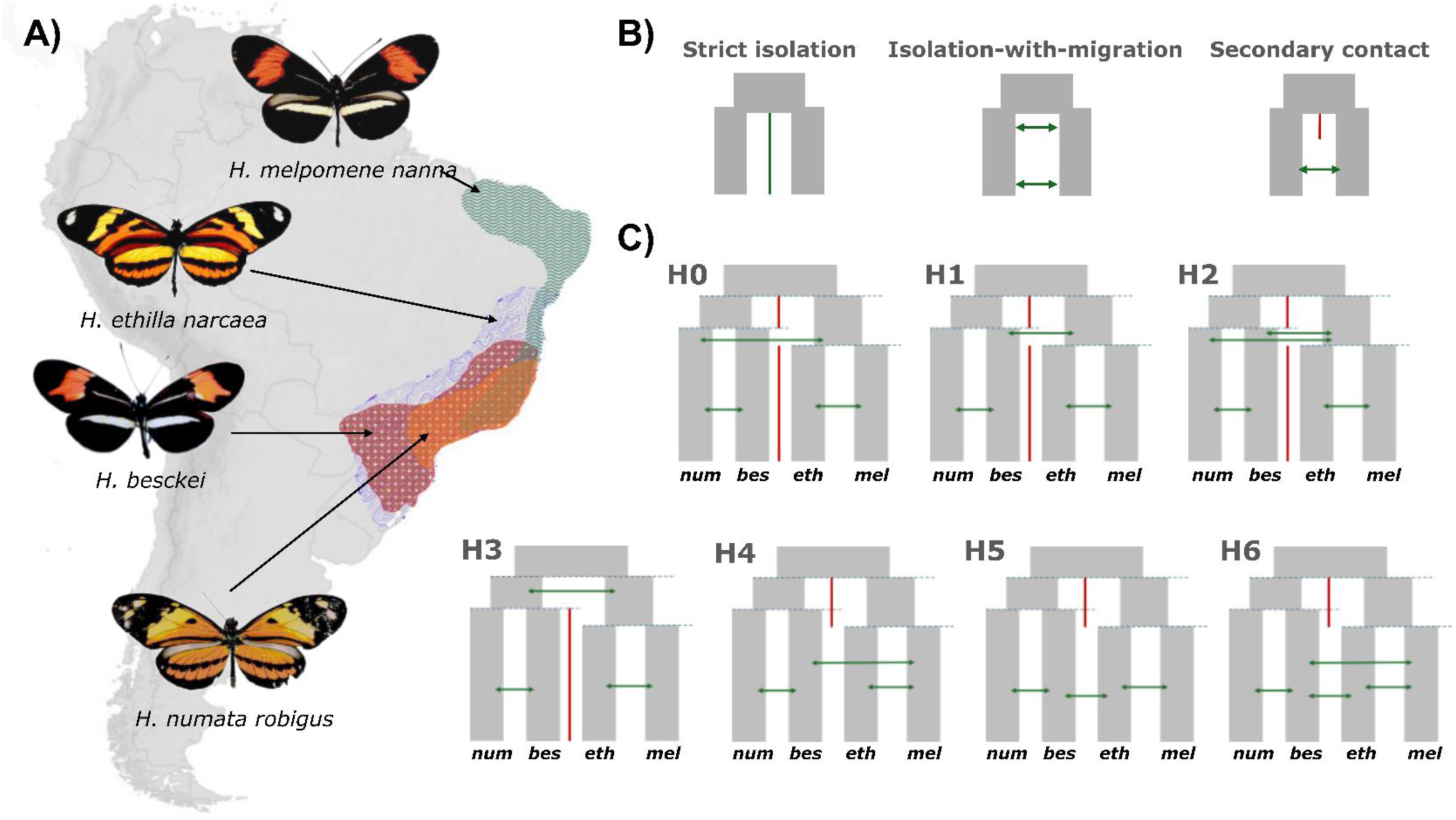
Study system and demographic models tested. (A) Schematic geographic distribution of the four focal Heliconius species. (B) Species pair scenarios tested with fastsimcoal2, differing on the occurrence and timing of gene flow. (C) Four-species scenarios testing the timing and strength of gene flow involving H. besckei. Hypotheses H0–H2 test whether gene flow between H. besckei and the melpomene–ethilla lineage can be explained indirectly by previously reported gene flow between H. numata and the melpomene–ethilla lineage, followed by gene flow between H. numata and H. besckei, or by direct gene flow between these lineages. We further tested if gene flow occurred before (H1–H2) or after (H4– H6) the divergence of H. melpomene and H. ethilla. Hypothesis H3 explains the current data by ancient gene flow between the ancestors of both lineages. Species names are simplified as follows: H. numata as num; H. besckei as bes; H. ethilla as eth; H. melpomene as mel. Butterfly images were obtained from the Butterflies of America website (Warren et al., 2024, http://www.butterfliesofamerica.com/).

Previous work has provided evidence that interspecific gene flow has shaped the evolutionary history of the *H. besckei* genome. Zhang et al. (2016) detected a signal of introgression between *H. besckei* and the ancestor of the *H. melpomene* lineage near the B/D mimicry region upstream of *optix*. Their analyses, however, found no similarly clear signal at the Yb locus, suggesting that introgression between these species was heterogeneous across the genome rather than uniformly associated with the shared phenotype. More broadly, introgression is widespread within the melpomene-silvaniform clade, including distantly related lineages (Thawornwattana et al. 2023). In the southeastern mountains of the Atlantic Forest, *H. besckei* also occurs in sympatry with *H. ethilla narcaea* (Figure 1), and putative phenotypic hybrids between the two taxa have been reported (Mallet et al. 2007, AVLF pers. obs, but see Brower 2018). However, these observations are based on phenotypic evidence, and the extent and persistence of gene flow between the two taxa, as well as the distribution of its signatures across the genome, remain unknown. Together, these observations raise the question of whether introgression involving *H. besckei* is concentrated in mimicry-associated genomic regions or instead reflects geographic contact and historical distribution changes.

Here, we investigate the evolutionary history of *H. besckei* by testing the extent and timing of interspecific gene flow in relation to mimicry, ecological context, and changes in geographic overlap. We distinguish two non-mutually exclusive scenarios: First, if selection for shared warning colouration promotes the persistence of introgressed mimicry alleles, we predict that *H. besckei* and the mimetic *H. melpomene nanna* comparisons will show elevated levels of introgression at the B/D region relative to comparisons involving non-mimetic sympatric and parapatric taxa. Second, if contemporary geographic contact is a stronger determinant of genome-wide introgression, we predict that *H. besckei* will show greater genome-wide allele sharing with the sympatric, non-mimetic *H. ethilla narcaea* than with the mimetic *H. melpomene nanna*, whose distributions overlap only narrowly. We further test whether Pleistocene climatic fluctuations repeatedly expanded and contracted suitable montane habitats for *H. besckei*. Periods of increased range size and connectivity should have created opportunities for secondary contact and gene flow involving *H. besckei* and lineages that are currently geographically separated. We therefore predict that demographic models allowing post-divergence migration will provide better support than models restricted to divergence without gene flow. Moreover, species distribution models should indicate shifts in the extent and connectivity of suitable habitats for *H. besckei*, with periods of greater connectivity expected during glacial periods, when drier and cooler climatic conditions may have favoured the expansion of habitats required for secondary contact in montane lineages. To test these predictions, we generated whole-genome sequence data from populations of *H. besckei* and related Atlantic Forest taxa and combined genome-wide analyses of introgression with demographic inference and species distribution modelling. This framework allows us to test whether locus-specific signals associated with mimicry can be distinguished from broader genome-wide patterns of gene flow associated with geographic contact.

## 2. Materials and Methods

### 2.1. Study system

We focused on four members of the silvaniform–melpomene radiation of *Heliconius* occurring in the Atlantic Forest, with *H. wallacei* included as an outgroup. Our focal species, *H. besckei*, is a montane Atlantic Forest species exhibiting a postman wing colour pattern. We also sampled the *postman* form *H. melpomene nanna*, which occurs mainly in lowland eastern Brazil. The distributions of *H. besckei* and *H. m. nanna* are broadly disjunct, although their ranges become locally parapatric in Espírito Santo (Fig. 1A; Brown & Freitas 2000). *Heliconius ethilla narcaea* is widely distributed throughout the Atlantic Forest and occurs across both lowland and montane habitats, overlapping with *H. besckei* (Figure 1A), and putative natural hybrids between the two taxa have been documented (Mallet et al. 2007) and observed in the field (AVLF pers. obs.). We therefore included *H. e. narcaea* to test whether contemporary geographic overlap is associated with gene flow involving *H. besckei*. Finally, we included *H. numata robigus*, an Atlantic Forest representative of the *H. numata* lineage and one of the closest extant relatives of *H. besckei* (Cicconardi et al. 2023; Thawornwattana et al. 2023). Together with *H. numata ethra*, *H. n. robigus* represents the species lineage in the Atlantic Forest (Brown Jr. 1976) and co-occurs with *H. besckei* on the southern coastal slopes of the Atlantic Forest (Brown & Freitas 2000; Francini et al. 2011). Unlike Amazonian populations of *H. numata*, Atlantic Forest populations retain the ancestral chromosomal arrangement and lack the inversion associated with the wing-pattern supergene polymorphism (Rodríguez de Cara et al. 2023).

### 2.2 Sample collection, DNA Extraction and Sequencing

We collected 47 specimens in the southeastern Brazilian Atlantic Forest, including *H. besckei, H. ethilla narcaea* and *H. melpomene nanna* (Table S1). We isolated DNA from either legs or thorax using the DNeasy Blood & Tissue Kit (Qiagen, Hilden, Germany) following the manufacturer’s protocol. Library preparation and whole-genome sequencing were performed by Novogene (UK) using an Illumina NovaSeq X Plus platform. We targeted an average sequencing depth of approximately 10-15X per sample. We further obtained whole-genome sequence data for 23 individuals from the NCBI Sequence Read Archive, corresponding to BioProjects PRJNA308754, PRJNA833307 and PRJEB8011 (Wallbank et al. 2016; Zhang et al. 2016; Rodríguez de Cara et al. 2023). The final dataset comprised 70 individuals: two *H. wallacei,* nine *H. numata robigus*, 15 *H. melpomene nanna*, 13 *H. besckei,* and 31 *H. ethilla narcaea* (Table S1).

### 2.3 Read processing, mapping and variant calling

We used *fastp* v.0.22.0 (Chen 2025) with default parameters to remove adapter sequences, poly-G tails, and bases with Phred quality scores < 15. Reads containing > 30% low-quality bases or shorter than 50 bp after trimming were excluded.

We mapped filtered reads to the *H. melpomene* v2.5 reference genome (Challi et al. 2016; Davey et al. 2016) using *bwa-mem2* v.2.2.1 (Vasimuddin et al. 2019) with the *MEM* algorithm and default options. We used *SAMtools* v1.14 (Danecek et al. 2021) to convert to BAM, sort and index our files, and *Picard Toolkit* v.2.27.5 (Broad Institute 2019) to remove PCR duplicates. Mapping quality, sequencing depth coverage, and GC content were assessed with *SAMtools*. The mean depth coverage among individuals sequenced for this study was 10.64X (Table S1).

We used *BCFtools* v.1.14 *mpileup* and *call* (Danecek et al. 2021) for variant and invariant sites calling on the 21 chromosomes. Site calling was performed separately for three datasets corresponding to the downstream analyses. The demographic dataset included *H. besckei*, *H. numata robigous*, *H. melpomene nanna* and *H. ethilla narcaea*. For introgression analyses, we generated two datasets containing the same *H. besckei* and *H. numata* individuals but differing in the third ingroup species: the melpomene dataset contained *H. melpomene nanna*, whereas the ethilla dataset contained *H. ethilla narcaea*. Both introgression datasets included the two *H. wallacei* samples as an outgroup. We filtered the resulting VCF files using *VCFtools* v.0.1.17 (Danecek et al., 2011), retaining biallelic sites with genotype quality ≥ 20 and a read depth of between 4 and 100 X. Sites with > 10% missing data across individuals, indels, and sites overlapping annotated repetitive regions of the *H. melpomene* reference genome (Lepbase; Challi et al. 2016) were excluded. For variable-site datasets, we additionally applied a minor allele frequency (MAF) ≥ 0.03. Additional filtering was applied where required for individual downstream analyses.

### 2.4 Species Differentiation and Nucleotide Diversity

To evaluate genomic differentiation among the four ingroup species, we ran a Principal Component Analysis (PCA) using Plink v2 (www.cog-genomics.org/plink/2.0/; Chang et al. 2015). PCA was performed on the variants-only demographic dataset after pruning variants for linkage disequilibrium using a 50-kb sliding window, a 10-kb step size and an r² threshold of 0.2.

Genomic differentiation between species was estimated using both relative and absolute measures of divergence. Genome-wide Weir and Cockerham *F_ST_* (1984) was calculated using *VCFtools* v.0.1.17 (Danecek et al. 2011). We calculated absolute genetic divergence (*Dxy*) in non-overlapping 10-kb windows with *pixy* (Korunes & Samuk 2021), which accounts for missing data and invariant sites. We estimated nucleotide diversity (**π**) and Watterson’s estimator of theta (*θ*) (Watterson 1975; Bailey et al. 2025) for each species in non-overlapping 10-kb windows using pixy and the all-sites demographic dataset.

### 2.5 Detection of introgression

To identify genomic regions potentially affected by introgression between *H. besckei* and *H. melpomene* (mimics) and between *H. besckei* and *H. ethilla* (sympatric non-mimics), we used two complementary approaches. First, in line with Zhang et al. (2016), we combined Patterson’s D statistic (Patterson et al. 2012) and the admixture proportion statistic *fd* with absolute divergence (*Dxy*) to distinguish candidate introgressed regions from patterns potentially generated by incomplete lineage sorting or shared ancestral polymorphism (Smith & Konforst 2013). Second, we used a topology-weighting framework (Martin & Belleghem 2017) implemented in twisst2 (Martin 2026) to identify genomic regions supporting genealogical relationships consistent with introgression.

#### 2.5.1 Patterson’s D, fd and Dxy

We used the script ‘*ABBABABAwindows.py*’ (Martin et al. 2015) to calculate Patterson’s *D* and *fd* in sliding windows of 5, 10, and 50 kb. We ran the analysis separately for the ethilla and melpomene datasets. We allowed up to 50% missing data per population and required a minimum SNP density of one SNP per 1,000 sites, following Zhang et al. (2016). We calculated the absolute divergence (*Dxy*) for the same genomic windows using pixy.

Because the B/D mimicry region upstream of *optix* has been identified as an introgressed region between *H. besckei* and *H. melpomene nanna* (Zhang et al. 2016), we used the signal at this region as an empirical reference for identifying genomic windows with unusually high allele sharing. We defined candidate windows using thresholds based on the values observed at the B/D region: *D* > 0.87 and *fd* > 0.75 for the 5- and 10-kb analyses and *fd* > 0.5 for the 50-kb analysis. To further reduce false positives, we retained candidate windows only when *Dxy* fell below the 25th percentile of the chromosome-specific *Dxy* distribution (Smith & Kronforst 2013; Martin et al. 2015). Candidate windows from the three window sizes were then merged using R v.4.3.2 (R Core Team, 2023) with the R package *GenomicRanges* (Lawrence et al. 2013). For each comparison, we quantified the number and total genomic extent of merged candidate regions and compared their genomic distribution between both datasets.

We further classified candidate regions according to whether they were detected in one or both comparisons (specific vs. shared). Because shared regions can arise from ancestral polymorphism or introgression predating the divergence of the relevant lineages, we interpreted this classification as evidence of differential allele sharing rather than as definitive evidence of independent introgression events.

#### 2.5.2 Topology weighting

As an independent assessment of introgression, we further investigated the different genealogical relationships along the genome with a topology weighting framework (Martin & Belleghem 2017) using twisst2 (Martin 2026). Local genealogies and genomic breakpoints were inferred using *sticcs* as implemented in twisst2 (Martin 2026). Because each dataset contained three species and one outgroup, only three rooted topologies were possible (Figure 2A).

**Figure 2.**
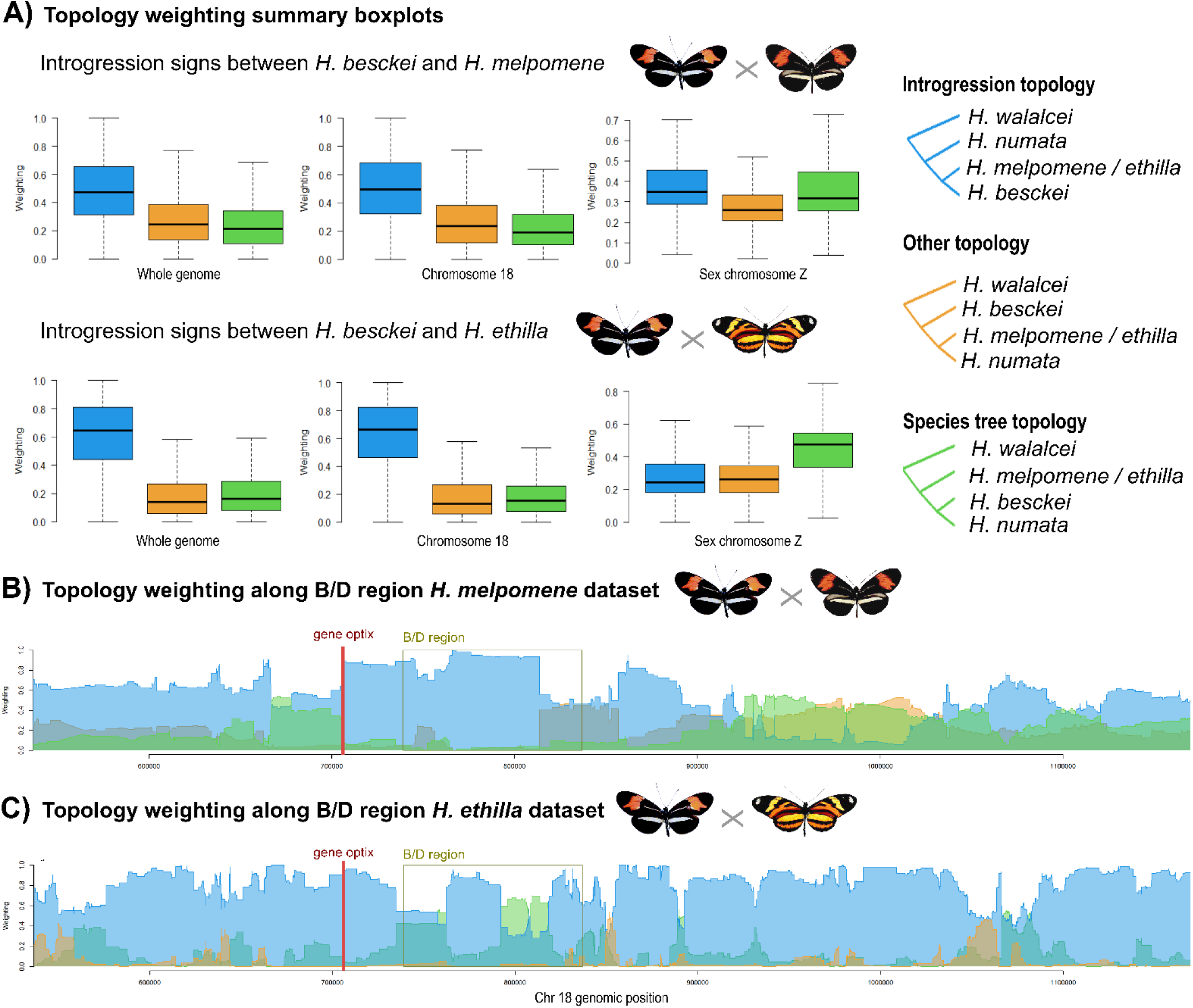
Signs of introgression between H. besckei and H. melpomene and between H. besckei and H. ethilla based on topology weighting results performed in Twisst2 for each dataset. (A) Summary boxplot of the weighting for each of the three possible topologies (right) for each dataset: the melpomene dataset (above) and the ethilla dataset (below). Colours correspond to the topologies shown on the right. (B) Topology weighting zoom around B/D regulatory region in chromosome 18 of the three possible topologies. Gene optix is highlighted in red, and the square upstream of it represents the B/D regulatory region. Results for both datasets are shown: the melpomene dataset (above) and the ethilla dataset (below). Colours correspond to the topologies shown on the right.

For each dataset, we identified genomic intervals in which the topology weight of the *H. besckei*-focal taxon topology exceeded 0.8. Contiguous intervals were merged, and the number, total size and genomic distribution of these sections were compared between datasets.

### 2.6 Demographic history

To assess whether *H. besckei* diverged in the presence or absence of gene flow, we used fastsimcoal2 v.2.8 (Excoffier et al. 2013, 2021; Marchi et al. 2024), which infers demographic parameters from the site frequency spectrum (SFS) using a coalescent simulation framework. Analyses were performed on the demographic dataset, including only data for the 20 autosomal chromosome scaffolds (Challi et al. 2016; Davey et al. 2016). We first performed species-pair analyses between *H. beskei* and one of the other focal species (Figure 1B), followed by more complex scenarios including the four species together (Figure 1C).

We used easySFS (Gutenkunst et al. 2009) to generate the input SFS files with projection values previously explored with the ‘*--preview*’ option (14, 18, 50, and 20 for *H. numata*, *H. besckei, H. ethilla, and H. melpomene,* respectively).

For each species pair, we tested three speciation scenarios: (1) strict isolation, assuming no gene flow after divergence; (2) isolation-with-migration, where continuous gene flow occurs after divergence; and (3) secondary contact, in which an initial period of isolation is followed by continuous gene flow (Figure 1B). We ran each scenario 100 times with 50 Expectation-Conditional Maximization (ECM) cycles (-L 50) and 1 million coalescent simulations (-n 1,000,000), with random seeds and a predefined parameter space for deme size (N: 1e^3^–2e^6^) and migration rate (m: 1e^-9^–0.1). Initial divergence times were set to 3,600,000–15,200,000 generations based on previous publications (Cicconardi et al. 2023; Thawornwattana et al. 2023), and we used a mutation rate of 2.9 x 10⁻⁹, estimated for *H. melpomene* (Keightley et al. 2014). To select the best run for each scenario, we retained the maximum estimated likelihoods and used the Akaike Information Criterion (AIC). Following model selection, to compare support among demographic models, we re-evaluated the composite likelihood of the best-fitting replicate of each scenario 100 times without parameter optimisation. Each evaluation was based on 1,000,000 coalescent simulations, generating a distribution of likelihood values for each model.

Based on the pairwise analyses, we designed six four-species demographic scenarios to test the timing and strength of gene flow between the *H. besckei* lineage and the *melpomene– ethilla* lineage (Figure 1C). Specifically, we asked whether the inferred gene flow between *H. besckei*, *H. ethilla narcaea* and *H. melpomene nanna* could be explained by the previously reported gene flow between *H. numata* and the *melpomene–ethilla* ancestral lineage (Thawornwattana et al. 2023), followed by more recent gene flow between *H. numata* and *H. besckei* (H0), or instead by direct gene flow between the *H. besckei* lineage and the *melpomene–ethilla* ancestral lineage. We further tested whether this direct gene flow occurred before the divergence of *H. ethilla* and *H. melpomene* (H1 and H2) or after their divergence (H4–H6).

In summary, H0 allowed for gene flow between *H. numata* and the ancestral lineage of *H. ethilla* and *H. melpomene*; H1 between *H. besckei* and the ancestral lineage of *H. ethilla* and *H. melpomene*; and H2 was a combination of H0 and H1, where gene flow between the ancestral lineages of *H. ethilla* and *H. melpomene* happened with both *H. numata* and *H. besckei*. H3 limited gene flow only between the two ancestral lineages (*H. besckei* – *H. numata* and *H. melpomene – H. ethilla*), while in H4 migration was allowed between *H. besckei* and *H. melpomene* and in H5 between *H. besckei* and *H. ethilla*; and finally, H6 allowed for gene flow between *H. besckei* and both *H. melpomene* and *H. ethilla*. Every scenario allowed for gene flow between *H. besckei* and *H. numata* and between *H. melpomene* and *H. ethilla* (Figure 1C).

We ran each scenario 50 times with 50 Expectation-Conditional Maximization (ECM) cycles (- L 50) and 500 K coalescent simulations (-n 500,000) with random seeds and the same predefined parameter space used for the species-pair scenarios. We constrained the divergence times of the species so that the divergence time matched previous estimates (Cicconardi et al., 2023), i.e., 3-4.8 Mya between the besckei-numata clade and the ethilla-melpomene clade; 2 Mya between *H. besckei* and *H. numata*; and at least 1.5 Mya between *H. ethilla* and *H. melpomene*. To select the best run for each scenario, we used AIC. In addition, we used the best-supported scenario to simulate 50 independent SFS datasets with fastsimcoal2 for parametric bootstrap, based on the parameter estimates from the best run. Each simulated SFS was then used as input to re-run the model with 50 ECM cycles and 500,000 coalescent simulations.

### 2.7 Species Distribution Models

We used species distribution models (SDMs) to estimate the climatic suitability of *H. besckei* under present and past climatic conditions. We compiled occurrence records using the R package *rgbif* (v.3.8.3) from GBIF (https://doi.org/10.15468/dl.mdte77; accessed 30 April 2026) and curated distributional datasets. To clean the records, we used *CoordinateCleaner* (v.3.0.1; Zizka et al., 2019) and *fuzzySim* (v.4.29; Vale & Brito, 2015), retaining only records with spatial uncertainty (<5 km) and collection years from 1970 onwards. Records with obvious geographic artefacts (e.g., country capitals, distribution points that lie at the geographic centre of the countries, institutional coordinates, GBIF headquarters, marine records, zero coordinates, and equal longitude–latitude values) were removed. Occurrences were additionally restricted to the known distribution of *H. besckei* in southeastern Brazil and spatially thinned at a minimum nearest-neighbour distance of 1 km to reduce sampling bias. We selected a 5 km uncertainty threshold after evaluating alternative cutoffs.

We obtained present-day climatic predictors from CHELSA bioclimatic variables (Karger et al., 2017) using the R package *geodata* v.0.6-6 (Hijmans et al. 2025). For spatial processing and raster handling, we used *terra* 1.7-83 (Hijmans, 2024) and *sf* 1.0-19 (Pebesma, 2018). Candidate climatic predictors were evaluated using pairwise correlation analyses and variance inflation factor (VIF) diagnostics implemented in *usdm* v.2.1-7 (Naimi 2023). The final models retained five climatic predictors based primarily on ecological relevance while considering collinearity among variables: temperature seasonality (BIO4), mean temperature of the warmest quarter (BIO10), precipitation of the wettest month (BIO13), precipitation of the driest month (BIO14), and precipitation seasonality (BIO15). These variables capture major thermal and precipitation gradients likely influencing the distribution of this montane Atlantic Forest specialist, including climatic stability, seasonal temperature constraints, and moisture availability during both humid and dry periods.

SDMs were fitted in R using Maxent implemented through the *maxnet* v.0.1.4 (Phillips, 2024) package. Models used linear, quadratic, and hinge features with a regularisation multiplier of 1.5. Background environments were represented by 10,000 randomly sampled points generated within a 300-km buffer around cleaned occurrence records. Models were subsequently projected beyond this calibration area across the broader American spatial extent covered by the environmental layers (125°W–25°W, 60°S–35°N). We assessed model performance using five-fold cross-validation and quantified it using the area under the receiver operating characteristic curve (AUC), and we calculated the true skill statistic (TSS) using *pROC* v.1.18.5 (Robin et al. 2011).

Then, we projected the models onto multiple palaeoclimatic scenarios from the PaleoClim database using *rpaleoclim* v.1.1.0 (Brown et al. 2018), including the Last Glacial Maximum (LGM, ∼21 Kya), MIS19 (∼787 Kya interglacial), and the M2 glacial event (∼3.3 Mya). The same set of climatic predictors (BIO4, BIO10, BIO13, BIO14, and BIO15) was used across all palaeoclimatic projections. To further investigate climatic dynamics during repeated Middle and Late Pleistocene cold phases associated with inferred secondary contact and introgression events, we generated additional palaeoclimatic projections using selected cold-stage time slices from the Oscillayers dataset (∼700 Kya, ∼450 Kya, and ∼140 Kya) (Gamisch, 2019).

We generated binary suitability maps using four alternative thresholds: the maximum TSS threshold, the 10th percentile training-presence threshold (p10), the 5th percentile training-presence threshold (p05), and the minimum training-presence threshold (minTP). The p10 threshold was used for the principal area estimates presented in the main text, while the alternative thresholds were used to assess the sensitivity of inferred palaeodistribution changes to threshold choice (Table S6). All palaeoclimatic projections used the same calibrated SDM and predictor framework to ensure comparability across time periods. We used *ggplot2* v.4.0.1 (Wickham et al. 2025) for figures and visualisations, *rnaturalearth* v.1.1.0 (Massicotte & South 2025) and *rnaturalearthdata* v.1.0.0 (South et al. 2024) for geographic map boundaries, and *vegan* v.2.6-6 (Oksanen et al. 2024) for spatial autocorrelation diagnostics.

## 3. Results

### 3.1 Strong species differentiation and a generally low nucleotide diversity

The PCA revealed clear overall genomic distinction among the four studied species, with no overlap between species clusters (Figure S1). Consistent with this, genome-wide estimates of genetic differentiation (*F*_ST_) and divergence (*Dxy*) clearly separated all species (Table S2). Pairwise *F_ST_* values – based on 1,891,222 polymorphic sites recovered for the demographic dataset – were high in all comparisons, ranging from 0.20 (*H. melpomene* – *H. ethilla narcaea*) to 0.74 (*H. numata – H. melpomene nanna*), with *Heliconius numata* consistently being the most differentiated species. Moreover, *H. numata* and *H. besckei*, despite being sister lineages, showed high *F_ST_* and *Dxy* values (0.72 and 0.005, respectively), while the sister lineages *H. ethilla* and *H. melpomene* showed lower values (0.20 and 0.002 for weighted *F_ST_* and *Dxy*, respectively; Table S2).

Genome-wide genetic diversity was low across all species (Table S3), as estimated by nucleotide diversity (**π**) and Watterson’s theta (*θ*). Among the four focal species, *Heliconius melpomene nanna* exhibited the lowest diversity (**π** = 0.00076, θ = 0.00215), whereas *H. numata* showed the highest (**π** = 0.00207, θ = 0.00397). In all species, **π** was consistently lower than θ, indicating an excess of low-frequency polymorphisms.

### 3.2 Evidence of pervasive introgression between *Heliconius* species

Both the combined *D* statistics, *fd and Dxy,* and the topology-weighting analysis revealed widespread genomic signatures of introgression between *H. besckei* and *H. melpomene nanna* (mimics), as well as between *H. besckei* and *H. ethilla narcaea* (sympatric).

Using the combined *D* statistics, *fd*, and *Dxy*, we found substantially more introgressed candidate windows between *H. besckei* and *H. ethilla narcaea* than between *H. besckei* and *H. melpomene nanna* (Figure 3). For 5 kb windows, 2.93% (7,456 windows) of the total number of windows that passed the filters supported introgression in the *H. ethilla narcaea* comparison, whereas it was 1.73% (4,307 windows) in the *H. melpomene nanna* comparison (Table S4). Similar patterns were observed for 10-kb (2.4% vs 1.6%) and 50-kb windows (3.2% vs 0.97%). Candidate introgressed regions, i.e., merged overlapping windows of the three different window sizes combined, were equally larger in the *H. ethilla narcaea* comparison, with mean sizes of 20.8 kb and 10.2 kb for *H. melpomene nanna.* We found 697 regions out of 1,753 that were specific to the dataset of *H. melpomene nanna and* 1,745 of a total of 2,442 to that of *H. ethilla narcaea* (Figure 3A).

**Figure 3.**
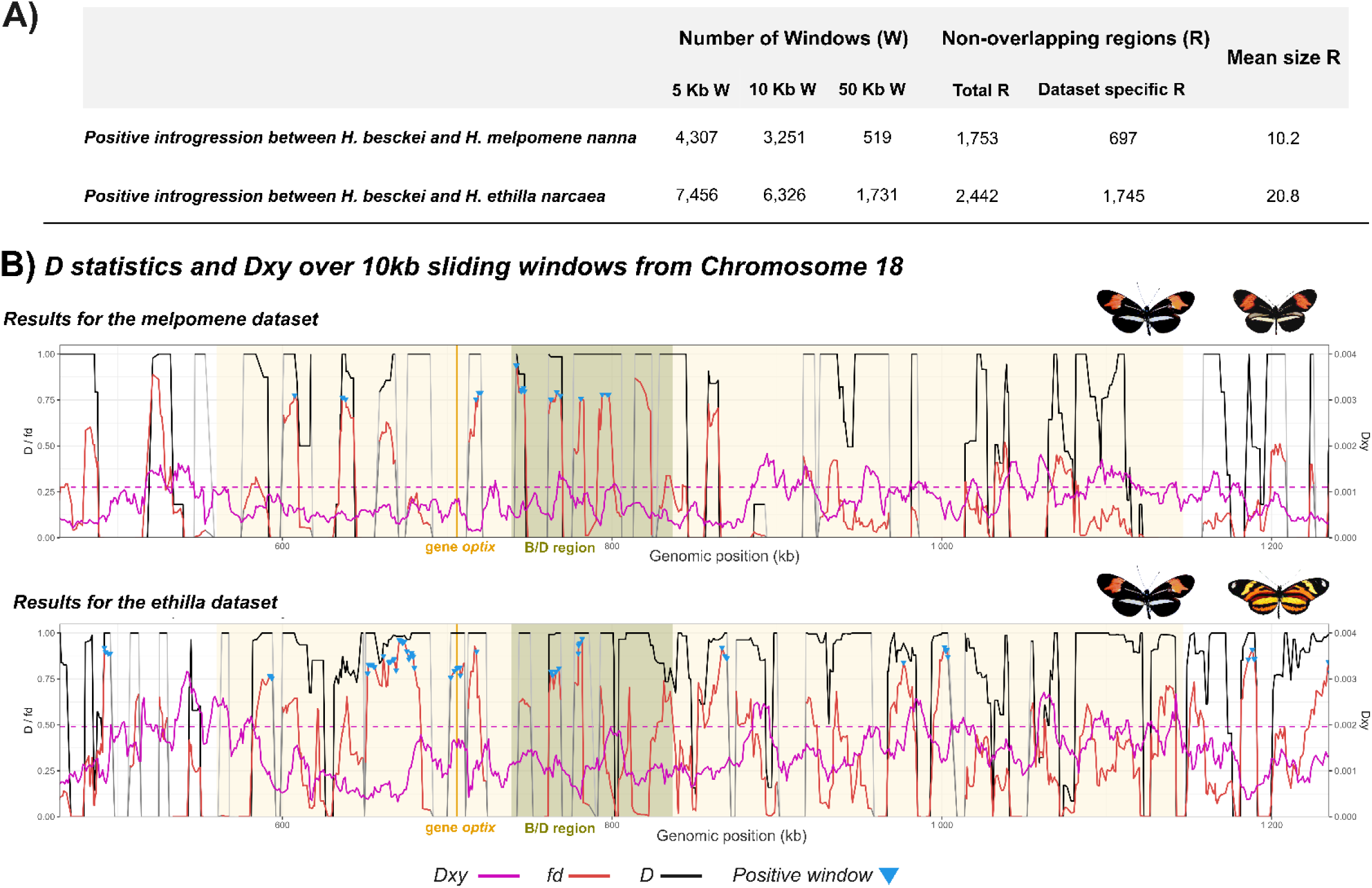
Signs of introgression between H. besckei and H. melpomene and between H. besckei and H. ethilla based on D-statistics results together with Dxy. (A) Genome-wide results for positive introgression windows and regions. A window was considered a positive sign of introgression based on D value (>0.87), fd (>0.75 for 5- and 10 kb windows and >0.5 for 50 kb windows), and Dxy (below the 25th percentile of the chromosome-specific distribution). The total number of positive windows (W) for the three different window size analyses, non-overlapping positive regions (R), and the mean size of the positive region among the two datasets are shown. (B) Chromosome 18 results around the B/D region (dark green shadow) that controls the postman red colour in Heliconius. Data for D statistics (black), fd (red) and Dxy (purple) of 10 kb sliding windows are shown. Windows that did not meet the filter criteria (fewer than 5 SNPs per 5 kb) and, therefore, were not used in the analysis, are represented here in grey for visualisation. Blue triangles point to windows that were found as a positive sign of introgression. Gene optix is represented with a yellow line. Results for both datasets are shown in the figure: for the melpomene dataset above and for the ethilla dataset at the bottom.

The topology-weighting analysis showed similar patterns. Across most of the genome, including chromosome 18 containing the B/D regulatory region, the topology grouping *H. besckei* with either *H. ethilla narcaea* or *H. melpomene nanna* received the highest support (Figure 2). In contrast, the species-tree topology was predominantly supported on the Z chromosome in the ethilla dataset. This provides further evidence of the high divergence of *H. numata* together with pervasive gene flow between *Heliconius* species (Figure 2).

Twisst2 also identified more genomic regions supporting introgression between *H. besckei* and *H. ethilla narcaea* than between *H. besckei* and *H. melpomene nanna*. A total of 11,091 sections (merged twisst2 intervals with the same higher-weighted topology) had topology weights > 0.8 consistent with introgression between *H. besckei* and *H. ethilla narcaea*, whereas 3,115 sections supported introgression with *H. melpomene nanna*, covering 32% of the genome compared to 11.26%, respectively. We found 9,772 out of a total of 14,206 sections to be dataset-specific: there were 8,602 sections uniquely supporting introgression between *H. besckei* and *H. ethilla narcaea,* compared with 1,170 sections uniquely supporting introgression between *H. besckei* and *H. melpomene nanna*, with a mean size of 6.7 kb in the former case and 4.3 kb in the latter.

### 3.3 Demographic modelling supports gene flow through *H. besckei*’s evolutionary history

For the species pair analyses, demographic inference with fastsimcoal2 consistently supported models including gene flow over strict isolation for all species pairs involving *H. besckei* (Table S5). The secondary contact scenario had the strongest support for all three pairwise comparisons based on AIC values (Table S5). However, for the comparison between *H. besckei* and *H. ethilla narcaea* and *H. besckei* and *H. numata*, the likelihood and AIC distributions of the secondary contact and isolation-with-migration scenarios largely overlapped, although ΔAIC between the two models was > 10 in both cases. Consequently, our data support gene flow with *H. besckei* after the speciation process but could not distinguish between a scenario of ancient secondary contact and one of continuous gene flow for these species pairs. For the four species models, the best supported scenario was H6, i.e., with gene flow between *H. besckei* and both *H. melpomene nanna* and *H. ethilla narcaea* after their split. The second and third best-supported scenarios also pointed to gene flow after the melpomene-ethilla divergence, probably being stronger between the lineage of *H. besckei* and *H. melpomene nanna* than with the *H. ethilla narcaea* lineage, since H4 showed a lower AIC than H5 (Table S6). The least likely scenario is the one with gene flow between the two ancestral populations of the melpomene-ethilla clade and the numata-besckei clade, H3.

Gene flow occurred bidirectionally in all models but was generally asymmetric in magnitude (Table S5; Table S6 and Figure 4b). In both the species-pair and the four-species scenario, the best-supported models’ migration was stronger from *H. beskei* into *H. numata* and from *H. melpomene* into *H. besckei*. In contrast, migration between *H. besckei* and *H. ethilla* was symmetric for the four-species model H6 but stronger from *H. besckei* to *H. ethilla* for the secondary contact scenario in the species-pair model. *H. besckei* also had a larger effective population size than *H. melpomene nanna* and *H. numata*, whereas *H. ethilla narcaea* exhibited by far the largest effective population size among the four analysed taxa (N=194,956 individuals by generation for the best run of H6). However, for the species pair scenarios, *H. numata* showed a slightly larger population size than *H. besckei*, as well as in the second-best model for the 4 species analyses (H4; Table S5 and Table S6). Furthermore, the last common ancestor showed the largest population, and the ancestor of the melpomene-ethilla clade was larger than the ancestor of the numata-besckei clade.

**Figure 4.**
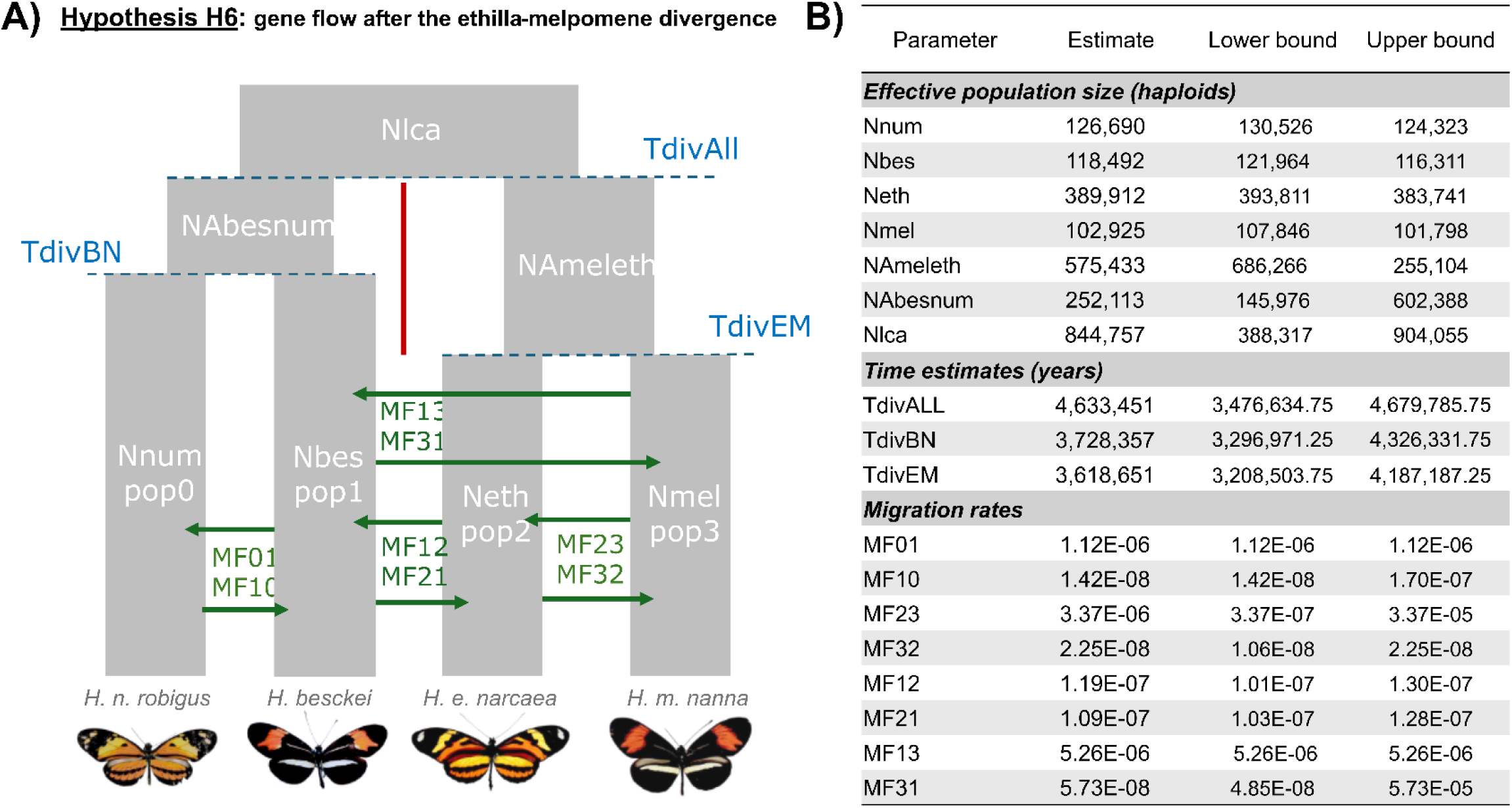
Demographic model and parameters estimated from the best-supported coalescent model (H6). (A) Schematic model tested with parameter names, which allowed for gene flow between H. besckei and both H. melpomene and H. ethilla after species divergence. (B) Demographic parameters estimated from the best run of the best-supported model (H6). Upper and lower bounds were estimated from 50 bootstrap replicates of the model using these point estimates to construct the simulated site frequency spectrum. The parameter names are defined in (A), and priors are specified in the input files reported on GitHub (https://github.com/RepullesM/Speciation-Heliconius-besckei.git). Butterfly images were obtained from the Butterflies of America website (Warren et al., 2024, http://www.butterfliesofamerica.com/).

### 3.4 Species distribution models indicate a larger suitable range for *besckei* during past glacial periods

After filtering and spatial thinning, 144 occurrence records were retained for model calibration (Figure 5A). The final model showed good predictive performance, with a mean five-fold cross-validation AUC of 0.957 and a mean TSS of 0.863, indicating high discrimination between suitable and unsuitable climatic conditions. The final model was based on five climatically meaningful predictors (BIO4, BIO10, BIO13, BIO14 and BIO15), selected according to their ecological relevance for tropical forest butterflies while accounting for multicollinearity. Correlation and variance inflation factor (VIF) diagnostics identified moderate correlations among several climatic variables, particularly between measures of temperature and precipitation seasonality. Nevertheless, all retained predictors showed acceptable VIF values and were retained because they represent distinct ecological processes likely influencing the distribution of *H. besckei*.

**Figure 5.**
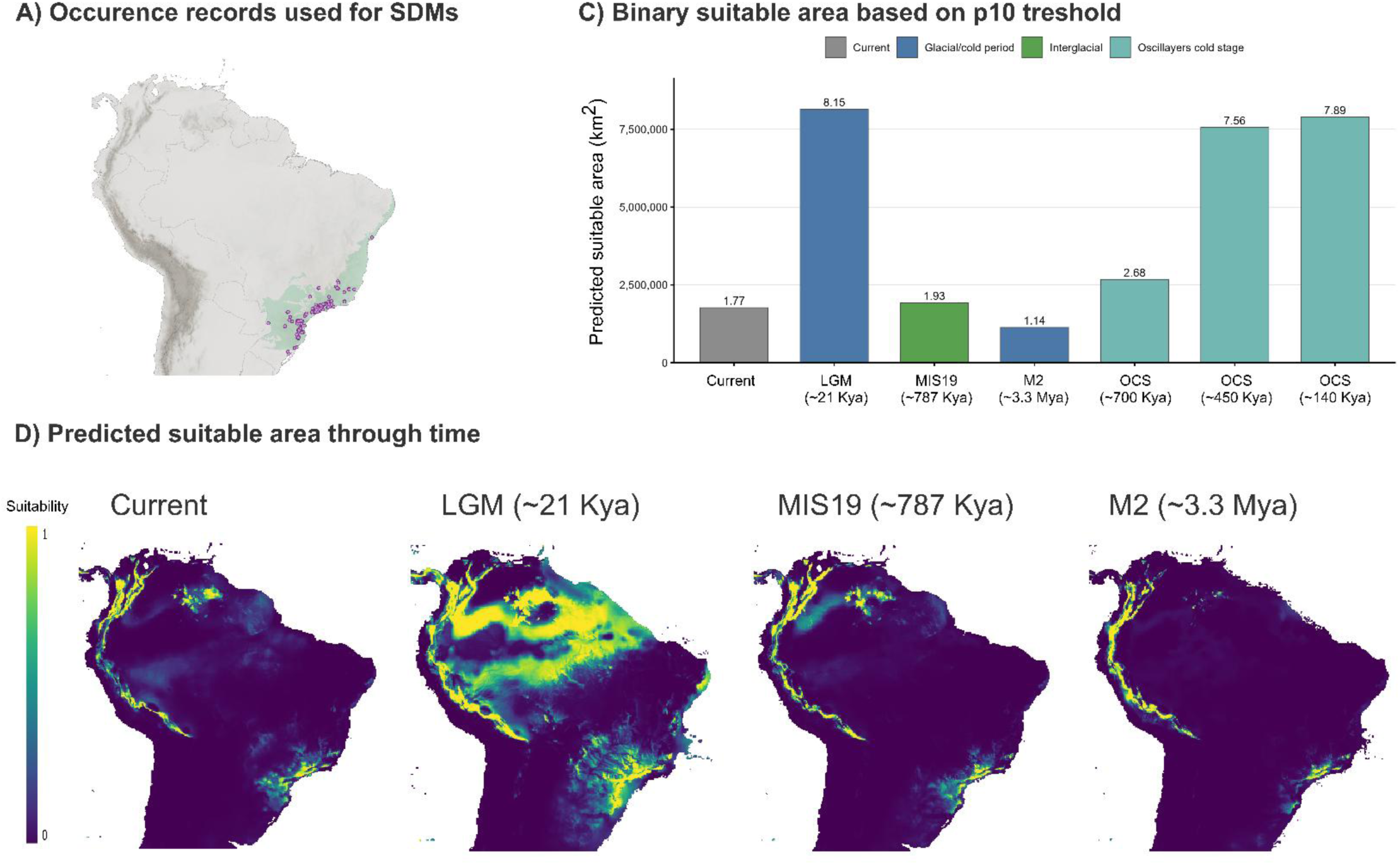
Predicted climatic suitability of Heliconius besckei under present-day and major palaeoclimatic scenarios. (A) Curated occurrence records used for Species Distribution Models (SDMs). The Atlantic Forest is shaded in green. (B) Predicted suitable area in km2 for the 10th percentile training-presence threshold (p10) for the different palaeoclimatic scenarios. (C) Predicted suitable area mapped for the Neotropical area for the p10 threshold results. The different panels show projections for the present (current), the Last Glacial Maximum (LGM; ∼21 Kya), the interglacial period MIS19 (∼787 Kya), and the M2 glacial event (∼3.3 Mya). Suitability was estimated using a MaxEnt (maxnet), species distribution model calibrated with five ecologically relevant climatic predictors (BIO4, BIO10, BIO13, BIO14 and BIO15). Colours indicate relative climatic suitability (0–1), with warmer colours representing higher suitability. Maps represent potential climatic suitability rather than confirmed historical occurrence.

The present-day projection closely matched the known distribution of *H. besckei* within the montane Atlantic Forest of southeastern Brazil (Figure 5), indicating that the model successfully captured the contemporary climatic niche of the species. Across all palaeoclimatic reconstructions, climatically suitable conditions remained largely restricted to the Atlantic Forest, and no climatically suitable corridor connecting the Atlantic Forest with Amazonian or Andean regions was recovered.

The Last Glacial Maximum (LGM; ∼21 Kya) was characterised by a marked expansion of climatically suitable area relative to the present (Table 2; Figure 5). Comparable expansions were consistently recovered for all three Oscillayers cold-stage reconstructions (∼700 Kya, ∼450 Kya and ∼140 Kya), with the largest suitable areas predicted during the ∼450 Kya and ∼140 Kya cold periods (Figure 5; Figure S2). In contrast, the M2 glacial event (∼3.3 Mya) produced a substantially smaller climatically suitable area than at present (Figure 5), whereas MIS19 (∼787 Kya) resulted in a distribution broadly comparable to the present (Figure 5). Although absolute estimates of suitable area differed among thresholding approaches, all methods consistently recovered repeated habitat expansions during Middle and Late Pleistocene cold periods (Table S15). This indicates that the inferred paleodistribution dynamics are robust to threshold selection rather than being driven by a particular binary classification criterion.

Overall, these reconstructions indicate repeated expansions of climatically suitable habitat during Middle and Late Pleistocene cold periods, after the divergence of the four lineages studied here. Importantly, however, all predicted expansions remained confined to the Atlantic Forest, providing no evidence for the formation of climatically suitable dispersal corridors towards Amazonian or Andean regions.

## 4. Discussion

*Heliconius besckei* is a geographically restricted Atlantic Forest butterfly that shares its *postman* wing pattern with the mostly parapatric *H. melpomene nanna*, while still hybridising with the sympatric, non-mimetic *H. ethilla narcaea* (Mallet et al. 2007). Our results reveal a complex history of post-divergence gene flow among these species. Demographic modelling provided evidence for gene flow between *H. besckei* and both *H. melpomene nanna* and *H. ethilla narcaea*, but genome-wide analyses showed contrasting genomic outcomes. Despite mimicry-related introgression reported in this and previous studies, the strongest genomic evidence for interspecific gene flow involves the sympatric, non-mimetic species. Recurrent contact between distinct species provides a sustained source of introgressed genetic variation, whose persistence is subsequently shaped by evolutionary processes, such as those underlying mimicry. Importantly, the widespread interspecific gene flow observed in this clade does not blur species boundaries, which remain genetically distinct.

The four *Heliconius* species studied are clearly distinct at the genomic level, with well-defined species clusters (Fig. S1) and substantial relative and absolute divergence among taxa, despite extensive evidence for gene flow. However, recurrent interspecific gene flow has shaped local genome genealogies, generating discordance between patterns of genomic differentiation and species relationships. *Heliconius numata robigus* was the most differentiated taxon, even from its sister lineage, *H. besckei.* Although *H. numata* forms natural hybrids with other *Heliconius* species in the Amazon forest (Mallet et al. 2007), Atlantic Forest populations seem to be genetically isolated. Rodriguez de Cara et al. (2023) already observed strong genetic differentiation and limited admixture between Atlantic Forest and Amazonian populations of *H. numata*, while our results further reveal that Atlantic Forest *H. numata* is highly differentiated from sympatric *Heliconius*, exceeding the differentiation observed among other species pairs. Indeed, *H. besckei* was less differentiated from *H. ethilla* and *H. melpomene* than from *H. numata*, despite being its sister lineage. This pattern may reflect the distinct demographic history of Atlantic Forest *H. numata* populations, differences in the extent of gene flow among the remaining lineages, or both.

The extensive gene flow identified here between *H. besckei* and the *melpomene–ethilla* clade likely homogenised portions of their genomes, generating widespread discordance between local genealogies and the species tree. Similar patterns have been reported in other *Heliconius* species, such as *H. pardalinus* and *H. elevatus*, where extensive gene flow has obscured genome-wide phylogenetic relationships (Rosser et al. 2024). Consistent with Thawornwattana et al. (2023), our topology-weighting analyses showed that the Z chromosome retains stronger support for the species tree than autosomal regions, suggesting that reduced introgression on sex chromosomes preserves phylogenetic signal. Therefore, genomic relationships among these species cannot be understood solely from their divergence history but also reflect subsequent episodes of interspecific gene flow, raising the question of when and under which demographic scenarios these exchanges occurred.

Our demographic model comparisons favoured scenarios involving gene flow after lineage divergence (Figure 4). Because recent phylogenomic reconstructions of *Heliconius* have detected ancestral introgression between *H. numata* and the melpomene-ethilla clade but not for *H. besckei* (Thawornwattana et al. 2023), our first hypothesis was that gene flow occurred first between *H. numata* and the melpomene-ethilla clade and subsequently between *H. numata* and *H. besckei* (H0, Figure 1C). However, we found better support for direct gene flow with *H. besckei* (H1, Figure 1C) than for a scenario mediated via *H. numata* (H0, Figure 1C). In addition, most of the shared variation found between *H. besckei* and the melpomene-ethilla clade is explained by gene flow after the *H. melpomene–H. ethilla* split (H4–H6 better supported than H0–H2, Figure 1C), approximately 3.6 Mya (Figure 4B). In line with this interpretation, window-based statistics (Figure 3) and topology weighting analysis recovered lineage-specific signals of introgression, suggesting that much of the genomic sharing is lineage-specific rather than ancestral.

Together with the demographic results, genome-wide introgression analyses suggest a strong geographical component to interspecific gene flow, with more widespread genomic exchange between sympatric species than comimics. Demographic models recovered bidirectional migration between *H. besckei* and the sympatric *H. ethilla*, but asymmetric migration between *H. besckei* and the mostly parapatric *H. melpomene nanna*. In addition, window-based and topology weighting analyses recovered substantially more introgressed regions and lower genome-wide differentiation (*F*_ST_) for the *H. besckei–H. ethilla* than for the *H. besckei–H. melpomene* comparisons (Figure 3A). Introgressed regions were also larger in the *H. besckei– H. ethilla* comparison, consistent with more recent or recurrent gene flow, as introgressed haplotypes are expected to become progressively fragmented by recombination and subsequent substitutions over time (Martin et al. 2013; Aguillon et al. 2022). However, this should be interpreted with caution, as the genomic extent of introgressed segments is also influenced by selection and demographic history (Aguillon et al. 2022).

We hypothesise that gene flow between *H. besckei* and *H. ethilla narcaea*, although potentially initiated earlier in their evolutionary history, persisted to the present facilitated by their contemporary sympatry, which might have favoured the occurrence of natural hybrids. In contrast, opportunities for contemporary gene flow between *H. besckei* and *H. melpomene nanna* are likely to be more restricted given their largely non-overlapping present-day distributions (Figure 1A; Brown 1979; Brown & Freitas 2000), as reflected by shorter and fewer regions showing evidence for introgression (Figure 3A and Table S4). In this case, selection associated with Müllerian mimicry may have favoured the retention of introgressed mimicry-associated alleles in *H. besckei*. Thus, the contrasting temporal and selective components of the two species pairs’ comparisons may reflect the complex evolutionary history of *H. besckei*: more geographically recurrent genome-wide exchange with *H. ethilla narcaea*, and more spatially restricted or selectively retained introgression from *H. melpomene nanna*.

The *H. besckei*-*H. melpomene* comparison provides an example of how selection may have facilitated the persistence of introgressed variation. We recovered a strong signature of introgression around the B/D locus between *H. besckei* and *H. melpomene nanna* (Figure 2B and Figure 3B), also consistent with the elevated migration rate from *H. melpomene* towards *H. besckei* in our demographic analysis results (Figure 4). In *Heliconius* butterflies, adaptive introgression at wing-pattern loci is driven by selection through Müllerian mimicry, where distinct species converge on similar phenotypes recognised by predators (Heliconius Genome Consortium 2012; Pardo-Díaz et al. 2012). In the *postman* phenotype, the red band of the pattern is mainly controlled by the B/D locus associated with the gene *optix* (Reed et al. 2011), and previous work suggested introgression from an *H. melpomene* ancestor to *H. besckei* (Zhang et al. 2016). However, based on our analysis, we cannot confirm or reject the direction of introgression for this region: the demographic analyses estimate genome-wide rather than locus-specific migration, while the tree topology- and *D-*statistic-based analyses cannot clearly distinguish adaptive from neutral introgression, nor do they indicate the direction of introgression (Martin et al. 2015). Moreover, the deep divergence between *H. besckei* and *H. numata* and the complex history of the silvaniform-melpomene clade complicate inference about the origin of the introgressed variant in *H. besckei*. In addition, introgression signals were also present genome-wide, not just at mimicry loci. While our analyses do not determine whether these regions were adaptively retained, studies in other *Heliconius* species have identified adaptive introgression involving loci associated with ecological and reproductive traits beyond mimicry (Montejo-Kovacevich et al. 2022; Rosser et al. 2024). Selectively retained introgressed variation outside mimicry loci may therefore also have contributed to the genomic landscape of *H. besckei*, although this remains to be tested. Given the largely non-overlapping present-day distributions of *H. besckei* and *H. melpomene nanna*, contemporary opportunities for gene exchange are likely more limited than with sympatric *H. ethilla*. The strongest genome-wide signal of gene flow occurs between the sympatric, non-mimetic species, whereas the clearest candidate adaptive signal occurs in the mimetic pair. Thus, the persistence of introgressed variation between *H. besckei* and *H. melpomene* may partly reflect the retention of advantageous alleles, while mechanisms such as assortative mating or selection against intermediate phenotypes may have shaped subsequent patterns of gene exchange. Whether gene exchange between *H. besckei* and the *H. melpomene* lineage occurred under sympatric or parapatric conditions, and how selection on mimicry loci contributed to the persistence of introgressed variation, requires further investigation.

Palaeodistribution models provide supporting geographic and temporal context for the inferred gene flow and demographic patterns. Climatically suitable areas for *H. besckei* were repeatedly larger during Middle and Late Pleistocene cold periods than at present (Figure 5), as also reported for other Atlantic Forest mountain species (Paz et al. 2019; da Silva Ribeiro et al. 2020). These expansions could have increased opportunities for secondary contact with currently separated or narrowly sympatric taxa, such as *H. melpomene nanna*. However, because climatic suitability was reconstructed only for *H. besckei*, these models identify periods of potential range expansion rather than directly demonstrating historical sympatry with donor lineages. The climatically suitable areas for *H. besckei* remained disconnected from Amazonian and Andean regions despite evidence for historical dispersal corridors for other taxa across the diagonal of open formations (Sobral-Souza et al. 2015; Ledo & Colli 2017). These findings suggest that the recurrent gene flow detected in *H. besckei*, including acquisition of mimicry-associated variation, likely occurred within the Atlantic Forest.

Overall, our results revealed recurrent interspecific gene flow in the Atlantic Forest endemic *H. besckei* despite substantial genomic differentiation from its congeners. Contrasting gene flow signatures from *H. melpomene nanna* and *H. ethilla narcaea* point to different temporal, geographic and selective contexts, while Pleistocene cold periods may have periodically increased opportunities for secondary contact. The contrasting genomic outcomes suggest that the extent of gene flow does not necessarily predict the persistence of particular introgressed variants. Together, these findings show that geographic restriction and strong species differentiation have not prevented recurrent genetic exchange during the diversification of *H. besckei* in the Atlantic Forest.

## Acknowledgements

We acknowledge computational resources provided by the e-INFRA CZ project (ID:90254), supported by the Ministry of Education, Youth and Sports of the Czech Republic. We thank the Instituto Chico Mendes de Conservação da Biodiversidade (ICMBio) for the collecting permits (10438-4 to 10438-8 and 10802-30 to 10802-40), including the permits for sampling threatened species (53016-10 to 53016-19). Access to genetic heritage was registered in the National System for the Management of Genetic Heritage and Associated Traditional Knowledge (SisGen) under the access code #AEAE189, in accordance with Brazilian Law No. 13,123/2015.

## Data Accessibility Statement

The raw data sequences generated in this study are available in NCBI assigned to the BioProject PRJNA1534588.

Bash and R scripts used for the genomic analyses are available in github: https://github.com/RepullesM/Speciation-Heliconius-besckei.git

## Benefit-Sharing Statement

Benefits Generated: A research collaboration was developed with scientists from the countries providing genetic samples, all collaborators are included as co-authors. The benefits of this research arise from the sharing of genomic data and research findings with the scientific community via public databases and publications. The study contributes to knowledge of the evolutionary history and biodiversity of the Brazilian Atlantic Forest.

## Author contributions

**MR**: Conceptualization (equal); Formal analysis (equal); Methodology (equal); Visualization (equal); Writing – original draft (equal). **TC**: Conceptualization (equal); Formal analysis (supporting); Methodology (equal); Supervision (equal); Writing – review and editing (supporting). **JK**: Formal analysis (equal); Methodology (supporting); Visualization (supporting); Writing – review and editing (equal). **KL**: Methodology (equal); Writing – review and editing (equal). **MJ**: Conceptualization (supporting); Methodology (supporting); Writing – review and editing (supporting). **NW**: Conceptualization (supporting); Supervision (supporting); Writing – review and editing (equal). **KLS:** Conceptualization (supporting); Writing – review and editing (supporting). **AF:** Conceptualization (equal); Funding acquisition (equal); Project administration (equal); Writing – review and editing (equal). **PMM:** Conceptualization (equal); Methodology (equal); Supervision (equal); Funding acquisition (equal); Project administration (equal); Writing – review and editing (equal).

## References

Abbott, R., Albach, D., Ansell, S., Arntzen, J. W., Baird, S. J. E., Bierne, N., Boughman, J., Brelsford, A., Buerkle, C. A., Buggs, R., Butlin, R. K., Dieckmann, U., Eroukhmanoff, F., Grill, A., Cahan, S. H., Hermansen, J. S., Hewitt, G., Hudson, A. G., Jiggins, C.,.. Zinner, D. (2013). Hybridization and speciation. Journal of Evolutionary Biology, 26(2), 229–246. 10.1111/j.1420-9101.2012.02599.x

Aguillon, S. M., Dodge, T. O., Preising, G. A., & Schumer, M. (2022). Introgression. Current Biology, 32(16), R865–R868. 10.1016/j.cub.2022.07.004

Bailey, N., Stevison, L., & Samuk, K. (2025). Correcting for Bias in Estimates of θw and Tajima’s D From Missing Data in Next-Generation Sequencing. Molecular Ecology Resources, 25(6), e14104. 10.1111/1755-0998.14104

Brower, A. V. Z. (2013). Introgression of wing pattern alleles and speciation via homoploid hybridization in Heliconius butterflies: A review of evidence from the genome. Proceedings of the Royal Society B: Biological Sciences, 280(1752), 20122302. 10.1098/rspb.2012.2302

Brower, A. V. Z. (2018). Alternative facts: A reconsideration of putatively natural interspecific hybrid specimens in the genus Heliconius (Lepidoptera: Nymphalidae). Zootaxa, 4499(1), 1–87. 10.11646/zootaxa.4499.1.1

Brown, J. L., Hill, D. J., Dolan, A. M., Carnaval, A. C., & Haywood, A. M. (2018). PaleoClim, high spatial resolution paleoclimate surfaces for global land areas. Scientific Data, 5(1), 180254. 10.1038/sdata.2018.254

Brown, K. S., & Mielke, O. H. H. (1972). The heliconians of Brazil (Lepidoptera: Nymphalidae). Part II. Introduction and general comments, with a supplementary revision of the tribe. Zoologica 57(1): 1–40. 10.5962/p.203234

Brown, K. S. (1976). An Illustrated Key to the Silvaniform Heliconius (Lepidoptera: Nymphalidae) with Descriptions of New Subspecies. Transactions of the American Entomological Society (1890-), 102(3), 373–484. JSTOR. http://www.jstor.org/stable/25078195

Brown, K. S. (1979). Ecologia geográfica e evolução nas florestas neotropicais. Zoology. Universidade Estadual de Campinas, Campinas, p. 256.

Brown, K. S. & Freitas, A.V.L. (2000). Diversidade de Lepidoptera em Santa Teresa, Espírito Santo. Boletim do Museu de Biologia Mello Leitão. 11(12), 71–116.

Capblancq, T., Després, L., Rioux, D., & Mavárez, J. (2015). Hybridization promotes speciation in *Coenonympha butterflies*. Molecular Ecology, 24, 6209–6222. 10.1111/mec.13479

Carnaval, A. C., Hickerson, M. J., Haddad, C. F. B., Rodrigues, M. T., & Moritz, C. (2009). Stability Predicts Genetic Diversity in the Brazilian Atlantic Forest Hotspot. Science, 323(5915), 785–789. 10.1126/science.1166955

Carnaval, A. C., & Moritz, C. (2008). Historical climate modelling predicts patterns of current biodiversity in the Brazilian Atlantic forest. Journal of Biogeography, 35(7), 1187– 1201. 10.1111/j.1365-2699.2007.01870.x

Challi, R. J., Kumar, S., Dasmahapatra, K. K., Jiggins, C. D., & Blaxter, M. (2016). Lepbase: The Lepidopteran genome database. bioRxiv, 056994. 10.1101/056994

Chang, C. C., Chow, C. C., Tellier, L. C., Vattikuti, S., Purcell, S. M., & Lee, J. J. (2015). Second-generation PLINK: rising to the challenge of larger and richer datasets. GigaScience, 4(1), s13742–015-0047–0048. 10.1186/s13742-015-0047-8

Chen, S. (2025). fastp 1.0: An ultra-fast all-round tool for FASTQ data quality control and preprocessing. iMeta, 4(5), e70078. 10.1002/imt2.70078

Cicconardi, F., Milanetti, E., Pinheiro de Castro, E. C., Mazo-Vargas, A., Van Belleghem, S. M., Ruggieri, A. A., Rastas, P., Hanly, J., Evans, E., Jiggins, C. D., Owen McMillan, W., Papa, R., Di Marino, D., Martin, A., & Montgomery, S. H. (2023). Evolutionary dynamics of genome size and content during the adaptive radiation of Heliconiini butterflies. Nature Communications, 14(1), 5620. 10.1038/s41467-023-41412-5

da Silva Ribeiro, T., Batalha-Filho, H., Silveira, L. F., Miyaki, C. Y., & Maldonado-Coelho, M. (2020). Life history and ecology might explain incongruent population structure in two co-distributed montane bird species of the Atlantic Forest. Molecular Phylogenetics and Evolution, 153, 106925. 10.1016/j.ympev.2020.106925

Danecek, P., Bonfield, J. K., Liddle, J., Marshall, J., Ohan, V., Pollard, M. O., Whitwham, A., Keane, T., McCarthy, S. A., Davies, R. M., & Li, H. (2021). Twelve years of SAMtools and BCFtools. GigaScience, 10(2), giab008. 10.1093/gigascience/giab008

The Heliconius Genome Consortium. (2012). Butterfly genome reveals promiscuous exchange of mimicry adaptations among species. Nature, 487(7405), 94–98. 10.1038/nature11041

Davey, J. W., Chouteau, M., Barker, S. L., Maroja, L., Baxter, S. W., Simpson, F., Merrill, R. M., Joron, M., Mallet, J., Dasmahapatra, K. K., & Jiggins, C. D. (2016). Major improvements to the Heliconius melpomene genome assembly used to Confirm 10 chromosome fusion events in 6 Million years of butterfly evolution. G3 Genes|Genomes|Genetics, 6(3), 695–708. 10.1534/g3.115.023655

Excoffier, L., Dupanloup, I., Huerta-Sánchez, E., Sousa, V. C., & Foll, M. (2013). Robust demographic inference from genomic and SNP data. PLOS Genetics, 9(10), e1003905. 10.1371/journal.pgen.1003905

Excoffier, L., Marchi, N., Marques, D. A., Matthey-Doret, R., Gouy, A., & Sousa, V. C. (2021). fastsimcoal2: Demographic inference under complex evolutionary scenarios. Bioinformatics, 37(24), 4882–4885. 10.1093/bioinformatics/btab468

Francini, R. B., Duarte, M., Mielke, O. H. H., Caldas, A., & Freitas, A. V. L. (2011). Butterflies (Lepidoptera, Papilionoidea and Hesperioidea) of the “Baixada Santista” region, coastal São Paulo, southeastern Brazil. Revista Brasileira de Entomologia, 55(1), 55–68.

Gamisch, A. (2019). Oscillayers: climatic oscillations reconstructed from terrestrial and marine fossil records for the past 5.4 million years. Scientific Data 6,254. 10.1038/s41597-019-0272-2

GBIF.org. (2026). GBIF Occurrence Download. 10.15468/dl.mdte77 (accessed 30 April 2026).

Gutenkunst, R. N., Hernandez, R. D., Williamson, S. H., & Bustamante, C. D. (2009). Inferring the joint demographic history of multiple populations from multidimensional SNP frequency data. PLOS Genetics, 5(10), e1000695. 10.1371/journal.pgen.1000695

Hijmans, R. J. (2024). terra: Spatial Data Analysis (Version 1.7-83) [R package]. Comprehensive R Archive Network (CRAN). https://CRAN.R-project.org/package=terra

Hijmans, R. J., Barbosa, M., Ghosh, A., & Mandel, A. (2025). geodata: Access Geographic Data (Version 0.6-6) [R package]. Comprehensive R Archive Network (CRAN). 10.32614/CRAN.package.geodata

Hines, H. M., Counterman, B. A., Papa, R., Moura, P. A., Cardoso, M. Z., Linares, M., Mallet, J., Reed, R. D., Jiggins, C. D., Kronforst, M. R., & McMillan, W. O. (2011). Wing patterning gene redefines the mimetic history of Heliconius butterflies. Proceedings of the National Academy of Sciences, 108(49), 19666–19671. 10.1073/pnas.1110096108

Jiggins, C. D. (2017). The Ecology and Evolution of Heliconius Butterflies. Oxford University Press. 10.1093/acprof:oso/9780199566570.001.0001

Joron, M., Frezal, L., Jones, R. T., Chamberlain, N. L., Lee, S. F., Haag, C. R., Whibley, A., Becuwe, M., Baxter, S. W., Ferguson, L., Wilkinson, P. A., Salazar, C., Davidson, C., Clark, R., Quail, M. A., Beasley, H., Glithero, R., Lloyd, C., Sims, S., … ffrench-Constant, R. H. (2011). Chromosomal rearrangements maintain a polymorphic supergene controlling butterfly mimicry. Nature, 477(7363), 203–206. 10.1038/nature10341

Karger, D. N., O. Conrad, J. Böhner, T. Kawohl, H. Kreft, R. W. Soria-Auza, N. E. Zimmermann, H. P. Linder, and M. Kessler. (2017). Climatologies at high resolution for the earth’s land surface areas. Scientific Data 4, 170122. 10.1038/sdata.2017.122

Keightley, P. D., Pinharanda, A., Ness, R. W., Simpson, F., Dasmahapatra, K. K., Mallet, J., Davey, J. W., & Jiggins, C. D. (2014). Estimation of the Spontaneous Mutation Rate in Heliconius melpomene. Molecular Biology and Evolution, 32(1), 239–243. 10.1093/molbev/msu302

Korunes, K. L., & Samuk, K. (2021). pixy: Unbiased estimation of nucleotide diversity and divergence in the presence of missing data. Molecular Ecology Resources, 21(4), 1359–1368. 10.1111/1755-0998.13326

Kozak, K. M., Joron, M., McMillan, W. O., & Jiggins, C. D. (2021). Rampant Genome-Wide Admixture across the Heliconius Radiation. Genome Biology and Evolution, 13(7), evab099. 10.1093/gbe/evab099

Kronforst, M. R., & Papa, R. (2015). The functional basis of wing patterning in Heliconius butterflies: The molecules behind mimicry. Genetics, 200(1), 1–19. 10.1534/genetics.114.172387

Lawrence, M., Huber, W., Pagès, H., Aboyoun, P., Carlson, M., Gentleman, R., Morgan, M. T., & Carey, V. J. (2013). Software for Computing and Annotating Genomic Ranges. PLOS Computational Biology, 9(8), e1003118. 10.1371/journal.pcbi.1003118

Ledo, R. M. D., & Colli, G. R. (2017). The historical connections between the Amazon and the Atlantic Forest revisited. Journal of Biogeography, 44(11), 2551–2563. 10.1111/jbi.13049

Leroy, T., & Heuertz, M. (2026). Adaptive Introgression in the Context of Climate Adaptation. Molecular Ecology, 35(8), e70333. 10.1111/mec.70333

Mallet, J. (2005). Hybridization as an invasion of the genome. Trends in Ecology & Evolution, 20(5), 229–237. 10.1016/j.tree.2005.02.010

Mallet, J., Beltrán, M., Neukirchen, W., & Linares, M. (2007). Natural hybridization in heliconiine butterflies: The species boundary as a continuum. BMC Evolutionary Biology, 7(1), 28. 10.1186/1471-2148-7-28

Mallet, J., & Gilbert Jr., L. E. (1995). Why are there so many mimicry rings? Correlations between habitat, behaviour and mimicry in Heliconius butterflies. Biological Journal of the Linnean Society, 55(2), 159–180. 10.1111/j.1095-8312.1995.tb01057.x

Marchi, N., Kapopoulou, A., & Excoffier, L. (2024). Demogenomic inference from spatially and temporally heterogeneous samples. Molecular Ecology Resources, 24(1), e13877. 10.1111/1755-0998.13877

Martin, S. H. (2026). A model-free method for genealogical inference without phasing and its application for topology weighting. Genetics, 232(1), iyaf181. 10.1093/genetics/iyaf181

Martin, S. H., Dasmahapatra, K. K., Nadeau, N. J., Salazar, C., Walters, J. R., Simpson, F., Blaxter, M., Manica, A., Mallet, J., & Jiggins, C. D. (2013). Genome-wide evidence for speciation with gene flow in Heliconius butterflies. Genome Research, 23(11), 1817– 1828. 10.1101/gr.159426.113

Martin, S. H., Davey, J. W., & Jiggins, C. D. (2015). Evaluating the Use of ABBA–BABA Statistics to Locate Introgressed Loci. Molecular Biology and Evolution, 32(1), 244– 257. 10.1093/molbev/msu269

Martin, S. H., & Van Belleghem, S. M. (2017). Exploring evolutionary relationships across the genome using topology weighting. Genetics, 206(1), 429–438. 10.1534/genetics.116.194720

Massicotte, P., & South, A. (2025). rnaturalearth: World Map Data from Natural Earth (Version 1.0.1) [R package]. Comprehensive R Archive Network (CRAN). https://CRAN.R-project.org/package=rnaturalearth

Mérot, C., Mavárez, J., Evin, A., Dasmahapatra, K. K., Mallet, J., Lamas, G., & Joron, M. (2013). Genetic differentiation without mimicry shift in a pair of hybridizing Heliconius species (Lepidoptera: Nymphalidae). Biological Journal of the Linnean Society, 109(4), 830–847. 10.1111/bij.12091

Merrill, R. M., Dasmahapatra, K. K., Davey, J. W., Dell’Aglio, D. D., Hanly, J. J., Huber, B., Jiggins, C. D., Joron, M., Kozak, K. M., Llaurens, V., Martin, S. H., Montgomery, S. H., Morris, J., Nadeau, N. J., Pinharanda, A. L., Rosser, N., Thompson, M. J., Vanjari, S., Wallbank, R. W. R., & Yu, Q. (2015). The diversification of Heliconius butterflies: What have we learned in 150 years? Journal of Evolutionary Biology, 28(8), 1417–1438. 10.1111/jeb.12672

Montejo-Kovacevich, G., Meier, J. I., Bacquet, C. N., Warren, I. A., Chan, Y. F., Kucka, M., Salazar, C., Rueda-M, N., Montgomery, S. H., McMillan, W. O., Kozak, K. M., Nadeau, N. J., Martin, S. H., & Jiggins, C. D. (2022). Repeated genetic adaptation to altitude in two tropical butterflies. Nature Communications, 13(1), 4676. 10.1038/s41467-022-32316-x

Moran, B. M., Payne, C., Langdon, Q., Powell, D. L., Brandvain, Y., & Schumer, M. (2021). The genomic consequences of hybridization. eLife, 10, e69016. 10.7554/eLife.69016

Nadeau, N. J., Pardo-Díaz, C., Whibley, A., Supple, M. A., Saenko, S. V., Wallbank, R. W. R., Wu, G. C., Maroja, L., Ferguson, L., Hanly, J. J., Hines, H., Salazar, C., Merrill, R. M., Dowling, A. J., ffrench-Constant, R. H., Llaurens, V., Joron, M., McMillan, W. O., & Jiggins, C. D. (2016). The gene cortex controls mimicry and crypsis in butterflies and moths. Nature, 534(7605), 106–110. 10.1038/nature17961

Naimi, B. (2023). usdm: Uncertainty Analysis for Species Distribution Models (Version 2.1-7) [R package]. Comprehensive R Archive Network (CRAN). 10.32614/CRAN.package.usdm

Oksanen, J., Simpson, G. L., Blanchet, F. G., Kindt, R., Legendre, P., Minchin, P. R., O’Hara, R. B., Solymos, P., Stevens, M. H. H., Szoecs, E., Wagner, H., Bedward, M., Bolker, B., Borcard, D., Carvalho, G., De Caceres, M., Durand, S., Evangelista, H. B. A., Hannigan, G., Hill, M. O., Lahti, L., Martino, C., Ouellette, M.-H., Ribeiro Cunha, E., Smith, T., Stier, A., Ter Braak, C. J. F., & Weedon, J. (2024). vegan: Community Ecology Package (Version 2.6-6) [R package]. Comprehensive R Archive Network (CRAN). 10.32614/CRAN.package.vegan

Pardo-Díaz, C., Salazar, C., Baxter, S. W., Merot, C., Figueiredo-Ready, W., Joron, M., McMillan, W. O., & Jiggins, C. D. (2012). Adaptive introgression across species boundaries in *Heliconius* butterflies. PLOS Genetics, 8(6), e1002752. 10.1371/journal.pgen.1002752

Patterson, N., Moorjani, P., Luo, Y., Mallick, S., Rohland, N., Zhan, Y., Genschoreck, T., Webster, T., & Reich, D. (2012). Ancient admixture in Human history. Genetics, 192(3), 1065–1093. 10.1534/genetics.112.145037

Paz, A., Spanos, Z., Brown, J. L., Lyra, M., Haddad, C., Rodrigues, M., & Carnaval, A. (2019). Phylogeography of Atlantic Forest glassfrogs (Vitreorana): When geography, climate dynamics and rivers matter. Heredity, 122(5), 545–557. 10.1038/s41437-018-0155-1

Pebesma, E. (2018). Simple Features for R: Standardized Support for Spatial Vector Data. The R Journal 10, 439–446. 10.32614/RJ-2018-009

Peres, E. A., Pinto-da-Rocha, R., Lohmann, L. G., Michelangeli, F. A., Miyaki, C. Y., & Carnaval, A. C. (2020). Patterns of Species and Lineage Diversity in the Atlantic Rainforest of Brazil. In V. Rull & A. C. Carnaval (Eds), Neotropical Diversification: Patterns and Processes (pp. 415–447). Springer International Publishing. 10.1007/978-3-030-31167-4_16 Broad Institute. (2019). Picard Toolkit [Computer software]. https://broadinstitute.github.io/picard/

Phillips, S. J. (2024). maxnet: Fitting “Maxent” Species Distribution Models with “glmnet” (Version 0.1.4) [R package]. Comprehensive R Archive Network (CRAN). https://CRAN.R-project.org/package=maxnet

R Core Team. (2023). R: A language and environment for statistical computing [Computer software]. R Foundation for Statistical Computing, Vienna, Austria. https://www.r-project.org/

Reed, R. D., Papa, R., Martin, A., Hines, H. M., Counterman, B. A., Pardo-Díaz, C., Jiggins, C. D., Chamberlain, N. L., Kronforst, M. R., Chen, R., Halder, G., Nijhout, H. F., & McMillan, W. O. (2011). Optix Drives the Repeated Convergent Evolution of Butterfly Wing Pattern Mimicry. Science, 333(6046), 1137–1141. 10.1126/science.1208227

Repullés, M., Chazot, N., Ramos, L. M.-, Espeland, M., Brandão, K. L. S.-, Antonelli, A., Freitas, A. V. L., & Matos-Maraví, P. (2026). Tracing the origins and evolution of nymphalid butterflies (Lepidoptera) in the Atlantic Forest. Ecography, 2026(6), e08419. 10.1002/ecog.08419

Robin, X., Turck, N., Hainard, A., Tiberti, N., Lisacek, F., Sanchez, J.-C., & Müller, M. (2011). pROC: an open-source package for R and S+ to analyze and compare ROC curves. BMC Bioinformatics, 12(1), 77. 10.1186/1471-2105-12-77

Rodríguez de Cara, M. Á., Jay, P., Rougemont, Q., Chouteau, M., Whibley, A., Huber, B., Piron-Prunier, F., Ramos, R. R., Freitas, A. V. L., Salazar, C., Silva-Brandão, K. L., Torres, T. T., & Joron, M. (2023). Balancing selection at a wing pattern locus is associated with major shifts in genome-wide patterns of diversity and gene flow. Peer Community Journal, 3. 10.24072/pcjournal.298

Rosser, N., Seixas, F., Queste, L. M., Cama, B., Mori-Pezo, R., Kryvokhyzha, D., Nelson, M., Waite-Hudson, R., Goringe, M., Costa, M., Elias, M., Mendes Eleres de Figueiredo, C., Freitas, A. V. L., Joron, M., Kozak, K., Lamas, G., Martins, A. R. P., McMillan, W. O., Ready, J., … Dasmahapatra, K. K. (2024). Hybrid speciation driven by multilocus introgression of ecological traits. Nature, 628(8009), 811–817. 10.1038/s41586-024-07263-w

Rosser, N., & Mallet, J. (2024). Interactive maps for visualizing geographic distributions and phenotypes. Tropical Lepidoptera Research, 34, 104–107. 10.5281/zenodo.13920055

Salazar, C., Baxter, S. W., Pardo-Díaz, C., Wu, G., Surridge, A., Linares, M., Bermingham, E., & Jiggins, C. D. (2010). Genetic evidence for hybrid trait speciation in Heliconius butterflies. PLOS Genetics, 6(4), e1000930. 10.1371/journal.pgen.1000930

Seehausen, O. (2004). Hybridization and adaptive radiation. Trends in Ecology & Evolution, 19(4), 198–207. 10.1016/j.tree.2004.01.003

Sheppard, P. M., Turner, J. R. G., Brown, K. S., Benson, W. W., & Singer, M. C. (1985). Genetics and the evolution of Mullerian mimicry in Heliconius butterflies. Philosophical Transactions of the Royal Society of London. Series B, Biological Sciences, 308(1137), 433–610. JSTOR. http://www.jstor.org/stable/2398716

Smith, J., & Kronforst, M. R. (2013). Do Heliconius butterfly species exchange mimicry alleles? Biology Letters, 9(4), 20130503. 10.1098/rsbl.2013.0503

Sobral-Souza, T., Lima-Ribeiro, M. S., & Solferini, V. N. (2015). Biogeography of Neotropical rainforests: Past connections between Amazon and Atlantic Forest detected by ecological niche modeling. Evolutionary Ecology, 29(5), 643–655. 10.1007/s10682-015-9780-9

South, A., Michael, S., Massicotte, P. (2024). rnaturalearthdata: World Vector Map Data from Natural Earth Used in ‘rnaturalearth’. (Version 1.0.0) [R package]. CRAN. https://CRAN.R-project.org/package=rnaturalearthdata

Thawornwattana, Y., Seixas, F., Yang, Z., & Mallet, J. (2023). Major patterns in the introgression history of Heliconius butterflies. eLife, 12, RP90656. 10.7554/eLife.90656

Vale, C. G., & Brito, M. P. (2015). Applying fuzzy logic to binary similarity indices in ecology. Methods in Ecology and Evolution 6, 853–858. 10.1111/2041-210X.12372

Vasimuddin, M., Misra, S., Li, H., & Aluru, S. (2019). Efficient architecture-aware acceleration of BWA-MEM for Multicore Systems. 2019 IEEE International Parallel and Distributed Processing Symposium (IPDPS), 314–324. 10.1109/IPDPS.2019.00041

Wallbank, R. W. R., Baxter, S. W., Pardo-Díaz, C., Hanly, J. J., Martin, S. H., Mallet, J., Dasmahapatra, K. K., Salazar, C., Joron, M., Nadeau, N., McMillan, W. O., & Jiggins, C. D. (2016). Evolutionary novelty in a butterfly wing pattern through enhancer shuffling. PLOS Biology, 14(1), e1002353. 10.1371/journal.pbio.1002353

Warren, A. D., Davis, K. J., Stangeland, E. M., Willmott, K. R. & Grishin, N. V. (2024). Illustrated Lists of American Butterflies. [9-III-2024]. http://www.butterfliesofamerica.com/

Watterson, G. A. (1975). On the number of segregating sites in genetical models without recombination. Theoretical Population Biology, 7(2), 256–276. 10.1016/0040-5809(75)90020-9

Weir, B. S., & Cockerham, C. C. (1984). Estimating F-Statistics for the analysis of population structure. Evolution, 38(6), 1358–1370. JSTOR. 10.2307/2408641

Wickham, H., Chang, W., Henry, L., Pedersen, T. L., Takahashi, K., Wilke, C., Woo, K., Yutani, H., Dunnington, D., van den Brand, T., & Posit, PBC. (2025). ggplot2: Create elegant data visualisations using the grammar of graphics (Version 4.0.1) [R package]. Comprehensive R Archive Network (CRAN). 10.32614/CRAN.package.ggplot2

Zhang, W., Dasmahapatra, K. K., Mallet, J., Moreira, G. R. P., & Kronforst, M. R. (2016). Genome-wide introgression among distantly related Heliconius butterfly species. Genome Biology, 17(1), 25. 10.1186/s13059-016-0889-0

Zizka, A., Silvestro, D., Andermann, T., Azevedo, J., Duarte Ritter, C., Edler, D., Farooq, H., Herdean, A., Ariza, M., Scharn, E., Svantesson, S., Wengström, N., Zizka, V., & Antonelli, A. (2019). CoordinateCleaner: standardized cleaning of occurrence records from biological collection databases. Methods in Ecology and Evolution 10, 744–751. 10.1111/2041-210X.13152

